# Short trains of transcutaneous vagus nerve stimulation increase online corticospinal excitability and pupil size in humans

**DOI:** 10.1101/2025.11.14.688461

**Authors:** Ronan Denyer, Shiyong Su, Mantosh Patnaik, Julie Duque

## Abstract

**Background:** Transcutaneous vagus nerve stimulation (tVNS) has emerged as a method for interrogating the role of the locus coeruleus (LC) norepinephrine system in human behavior. Tuning of excitability in the corticospinal tract is central to many cognitive and motor processes, but little is known about how the LC contributes to this tuning. In particular, no existing studies have examined the effect of tVNS on corticospinal excitability “online” during active stimulation, where the largest effects on pupil size are observed.

**Method:** To address this question, we delivered repeated 4-second trains of tVNS and sham stimulation and elicited MEPs during stimulation trains (online) and shortly after train offset (offline). Pupil size was concurrently recorded throughout each train.

**Results:** We discovered that tVNS significantly increases corticospinal excitability compared to sham stimulation, but only when measured online and not offline. The excitatory effects on corticospinal excitability were greater in the latter half of tVNS trains. Pupil size was also significantly increased by tVNS compared to sham; however, the effect on pupil size peaked earlier during the tVNS trains compared to corticospinal excitability. In line with these distinct temporal profiles, changes in corticospinal excitability and pupil size were not significantly correlated, likely reflecting differences in the anatomical circuits underpinning each effect.

**Conclusions:** This work demonstrates for the first time that tVNS increases corticospinal excitability at rest, but the effect only emerges when corticospinal excitability is measured online during active tVNS. Implications for basic and clinical neuroscientific research are discussed.

## 1 Introduction

Transcutaneous vagus nerve stimulation (tVNS) is an emerging form of non-invasive brain stimulation that shows promise as a method to better understand the role of the locus coeruleus (LC) in human behavior and neurological disease. tVNS involves electrical stimulation of the cymba conchae within the ear, which drives activation of the auricular branch of the vagus nerve^1–3^. This activation spreads to the nucleus tractus solitarii, which in turn activates the LC^4–6^ and releases norepinephrine (NE) throughout the central nervous system (CNS). The effect of tVNS on LC activity is evidenced through studies showing that tVNS increases neurophysiological markers of NE release, including pupil size^7–10^, heart rate^11^, and skin conductance^12^. Functional magnetic resonance imaging further indicates that LC activity is increased during tVNS. Similarly, animal studies show dose-dependent increases in LC activity during VNS and increases in basal LC firing found after repeated bouts of VNS. These induced increases in LC activity are associated with excitation across a broad array of cortical and subcortical regions^13–15^.

While the effect of tVNS on traditional makers of NE release are relatively well understood, little is known about its effect on corticospinal excitability. This is important to address since the corticospinal tract is the largest descending pathway in the brain and critical for voluntary motor control. Excitability in the corticospinal tract can be indexed with high temporal precision by analyzing the amplitude of motor evoked potentials (MEPs) in peripheral muscle(s) elicited by single pulses of transcranial magnetic stimulation (TMS) over the primary motor cortex (M1). Studies combining TMS with behavioral tasks have shown that rapid fluctuations in corticospinal excitability are a feature of multiple cognitive and motor processes^16–18^, including motor preparation^19–32^, decision making^33,34^, movement cancellation^35–38^, and complex bimanual control^39,40^. The prevalence of corticospinal tract disruption in numerous motor system diseases including stroke^41–44^, multiple sclerosis^45^ and Parkinson’s disease^46,47^ further showcases the centrality of the corticospinal activity in adaptive movement.

Existing investigations into the effect of tVNS on corticospinal excitability have yielded mixed results. All studies have focused on the “offline” changes in corticospinal excitability before and after a 30-60 minute bout of continuous tVNS, with the majority finding no significant effect on corticospinal excitability^48–50^. Other work showed an increase in corticospinal excitability after tVNS^51^, but this was with electrodes placed bilaterally on the neck, making it difficult to ascertain the locus of stimulation.

The lack of a clear effect of tVNS on corticospinal excitability may be caused in part by the focus on measuring corticospinal excitability “offline” before and after a bout of tVNS. In these scenarios, even if tVNS did influence activity in the corticospinal tract, it is possible that homeostatic plasticity may have been engaged to normalize cortical activity, resulting in no observable changes after the stimulation^52–54^. A more appropriate approach might be to assess effects on corticospinal excitability “online” while tVNS is actively applied. Work from our lab^7^ and others^8,9^ indicates that tVNS has very rapid acting effects on pupil size, with the application of 4-second trains of tVNS at rest resulting in a significant increase in pupil size often less than a second after tVNS onset when compared to a sham stimulation condition^7^. Notably, pupil size returned to baseline levels just 1 second after tVNS offset^7^.

To assess the online effects of tVNS on corticospinal excitability, we elicited MEPs before (baseline), during (online) and after (offline) repeated 4-second trains of tVNS. These MEP data were compared with MEPs elicited during intensity matched sham stimulation trials in which 4-second trains of stimulation were applied to the ear lobes. We predicted that corticospinal excitability would be increased by tVNS, but only when measured online and not offline. We concurrently recorded pupil size throughout every tVNS/sham train to assess the effect of tVNS on this traditional marker of NE release. We predicted that a significant increase in tVNS relative to sham would be found^7^. Finally, we also predicted that normalized increases in corticospinal excitability and pupil size would be correlated.

## 2 Methods

### 2.1 Participants and experimental setup

Twenty-nine participants were recruited; one was excluded due to a technical issue with tVNS, leaving 28 (21 female/7 male; age 25.1 ± 3.2; 26 right-handed/2 left-handed). All had normal or corrected vision, no neurological or psychiatric disorders, and no physical injuries. Participants provided written informed consent, received financial compensation, and the study was approved by the Ethics Committee of the Université catholique de Louvain and the Cliniques universitaires Saint-Luc (Brussels, Belgium; 2024/05JUI/292 - NEURAL OSCILLATIONS) in accordance with the Declaration of Helsinki. Experiments were conducted in a quiet, moderately lit room. Participants sat at a chin rest with hands flat on a table, facing a computer screen (**Figure 1A**).

**Figure 1.**
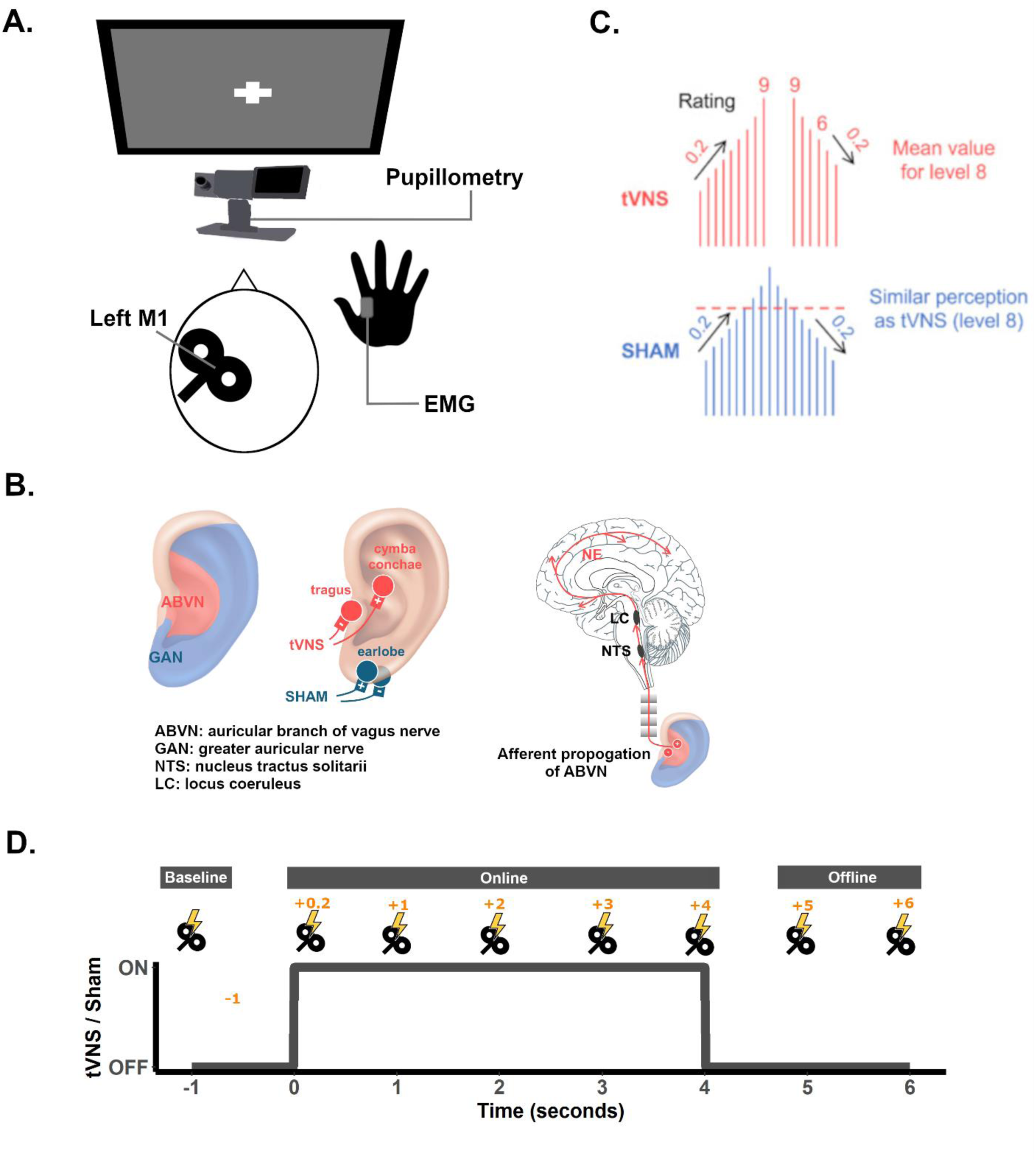
Experimental setup and design. **A**. Participants sat in front of a monitor and fixated on a white cross displayed in the center of a black screen while receiving tVNS/SHAM stimulation every 14–15s at rest. TMS was deliver over left M1 to elicit motor evoked potentials (MEPs) in the right flexor dorsal interosseous (FDI). An eye-tracking camera was used to capture pupillometry and EMG was used to capture right FDI MEPs. **B.** Electrode montage in tVNS and SHAM conditions. Electrodes were placed on the cymba conchae and tragus of the left ear in tVNS condition^7,55–57^. This generates an electric field covering the area innervated by the auricular branch of the vagus nerve (ABVN)^1–3^, thereby specifically activating the vagal afferent pathway up to the nucleus tractus solitarii (NTS), which in turn transmits impulses to locus coeruleus (LC)^4–6^ and activates norepinephrine (NE) neurons that project throughout the CNS. For sham stimulation, electrodes were put on the left earlobe. This allows to generate an electric field covering the area of the greater auricular nerve (GAN), where stimulation is not expected to induce brainstem activation^8^. Pulses (200 μs) were delivered at 25 Hz for 4 s in both conditions. **C.** A “Method of Limits” procedure was used to select the maximal comfortable tVNS intensity for each participant. Participants received increasing and decreasing series of 4s tVNS trials and rated their subjective sensation for each stimulation on a 10-point scale. The intensity started at 0.1 mA and was increased by 0.2 mA until participants reported a sensation of 9. Before starting the decreasing series, the same intensity was repeated. Then, it was reduced trial by trial in 0.2 mA steps until a subjective sensation of 6 or below was experienced. The final intensity for tVNS was calculated based on the average of the intensities rated as 8. To match the perception in the two conditions, intensity in the sham condition was set to the equal perception of the final intensity in tVNS condition. **D.** Participants received repeated and randomly interleaved 4-second trains of tVNS and sham stimulation. MEPs were elicited at one of 8 timepoints relative to the start of tVNS/sham trains. Trains of tVNS/sham were separated by a 9-10 second intertrial period.

### 2.2 tVNS

tVNS and sham stimulation were delivered in 4-s trains via a custom electrical stimulation platform^7^. Stimulation targeted the left cymba conchae (tVNS) or left earlobe (sham) using flexible pad electrodes (Axiothera, Germany) connected to Digitimer DS7A devices (Digitimer, UK), with intensity and pulse duration controlled on the DS7A. Both devices were linked to a Master-9 multichannel stimulator (A.M.P.I., Israel) to control pulse frequency and train duration. Electrode placement mirrored previous work (**Figure 1B**), and intensity was calibrated according to established protocols (**Figure 1C**).

### 2.3 TMS

Single-pulse TMS and surface electromyography (EMG) were used to determine resting motor threshold (RMT) and measure corticospinal excitability. EMG was recorded from the right flexor dorsal interosseous (FDI) muscle using bipolar Neuroline electrodes (Ambu, Denmark) with a ground on the ulnar styloid process. Signals were amplified ×1,000, bandpass filtered (50–450 Hz; Digitimer, UK) and sampled at 2,000 Hz using Signal (Cambridge Electronic Design, UK). TMS was delivered over left M1 using a MagVenture stimulator (Denmark) with a C-B60 coil, guided by a Visor2 neuronavigation system (ANT Neuro, Germany). The “hotspot” which elicited consistent MEPs in the contralateral FDI was identified. RMT was then delineated by finding the lowest stimulation intensity that elicited ≥0.05 mV MEPs in 5 out of 10 trials. TMS was applied at an intensity of 115% RMT throughout the experiment.

### 2.4 Combined tVNS/TMS/Pupillometry

Participants completed 10 blocks of 40 stimulation trains each (20 tVNS, 20 sham) while at rest. Single TMS pulses were applied at −1, +0.2, +1, +2, +3, +4, +5, or +6 s relative to train onset (**Figure 1D**) to assess corticospinal excitability, measured via peak-to-peak MEP amplitude. Two MEPs were elicited per timepoint per condition, totaling 36 MEPs per block and 20 MEPs per trial type across the experiment. Each block also included two tVNS and two sham trains without TMS to assess pupil responses in the absence of TMS pulses. Train types were randomly interleaved, separated by 9–10 s intertrial intervals to allow pupil time to return to baseline^7^. Pupillometry was recorded continuously at 1000 Hz using an Eyelink 1000+ eye tracker (SR Research, Canada), and train/TMS timing was controlled using a custom MATLAB script. Participants completed a brief Eyelink calibration procedure before TMS blocks to ensure stable pupil size measurements.

### 2.5 Data processing

#### 2.5.1 TMS/EMG data

TMS/EMG data were analyzed offline using the VETA MATLAB toolbox. MEP peak-to-peak amplitude was the primary variable. MEPs were automatically detected with ‘findEMG.m’ and visually verified with ‘visualizeEMG.m’; onsets and offsets were adjusted as needed. Trials showing >0.05 mV background EMG within 50 ms pre-TMS were excluded (∼2%). MEPs were normalized to each participant’s mean baseline amplitude (1 s pre-tVNS/sham onset) and expressed as ratios. Extreme outliers were removed before averaging to minimize skew.

#### 2.5.2 Pupillometric data

Pupil data were analyzed offline using custom MATLAB scripts. One participant was excluded due to poor tracking. Blink periods were removed via linear interpolation (±100 ms) and data low-pass filtered at 10 Hz (fourth-order Butterworth, zero-phase). Trials with >50% interpolated samples were excluded (∼3%). Data were segmented from 1 s pre- to 12 s post-event for tVNS/sham trains and 5 s post-event for TMS pulses. Each trial was baseline-corrected using the mean pupil size in the 1 s pre-event period to control for session-wide fluctuations.

### 2.6 Data analysis

#### 2.6.1 Model selection

For most analyses, generalized linear mixed-effects regression (GLMER) with a gamma distribution and identity link was used. This approach accommodates the skewness typical of neurophysiological data^58,59^ and provides superior handling of interindividual variability through random effects compared to traditional ANOVA or linear regression^60–62^. Participant ID was always included as a random intercept. Simple main effects were tested using estimated marginal means with Tukey-adjusted pairwise comparisons.

#### 2.6.2 Effect of tVNS on corticospinal excitability

This study primarily assessed whether tVNS modulates corticospinal excitability when applied *online* (during stimulation) versus *offline* (after stimulation). MEP data were divided into three timeframes: Baseline (−1 s pre-train), Online (+0.2–4 s post-onset), and Offline (+5–6 s). Baseline MEPs, collected prior to the onset of either stimulation type, were entered as identical values for both tVNS and sham conditions to provide a shared reference level for the GLMER, allowing interaction terms to reflect stimulation-specific deviations. Normalized MEPs were averaged within each timeframe and stimulation condition, and a GLMER tested the fixed effects of Time (Baseline, Online, Offline) and Stimulation Type (tVNS, Sham).

A secondary analysis examined whether tVNS effects on corticospinal excitability were uniform across all online timepoints or specific to certain intervals. Mean normalized MEPs were computed at each timepoint (−1 [shared reference], +0.2, +1, +2, +3, +4 s), and a GLMER assessed fixed effects of Time and Stimulation Type on normalized MEP amplitude.

#### 2.6.3 Effect of tVNS on pupil size

To assess pupil responses to 4-s tVNS trains, only no-TMS trials were analyzed, to control for the potential contaminating effects of TMS pulses. Baseline-normalized pupil data were averaged across 40 trials per stimulation condition. Differences between tVNS and sham were tested using a cluster-based paired *t*-test (20,000 Monte Carlo permutations). Mean pupil size within significant clusters was entered into a GLMER with Stimulation Type as a fixed factor.

#### 2.6.4 Association between corticospinal excitability and pupil size

An exploratory analysis tested whether changes in corticospinal excitability correlated with changes in pupil size. For each post-onset timepoint (+0.2–6 s), MEP change scores (tVNS − sham) and pupil change scores (tVNS − sham, 500 ms pre-TMS window) were computed. An LMER assessed Pupil Change and Time as fixed effects on MEP Change.

#### 2.6.5 Effect of TMS pulses on pupil size

To isolate TMS effects on pupil size, only post-train sham trials were analyzed, with pupil data from no-TMS trials used as control to compare changes against. Baseline-normalized pupil data (−1 to +5 s around TMS onset time) were averaged across 40 trials per condition. A cluster-based paired *t*-test (20,000 permutations) compared TMS and no-TMS conditions, and mean pupil size from significant clusters was entered into a GLMER with TMS Type as a fixed factor.

## 3 Results

### 3.1 tVNS trains drive online but not offline increases in corticospinal excitability

A GLMER testing whether tVNS modulated online corticospinal excitability showed a significant Time × Stimulation Type interaction (β = 0.05, 95% CI [0.03, 0.08], *p* < 0.001). Estimated marginal means indicated greater excitability online during tVNS than sham (μ = 0.051 ± 0.014, *p* < 0.0005), but no difference offline after stimulation (*p* = 0.28; **Figure 2A**), suggesting effects were confined to the active stimulation period.

**Figure 2.**
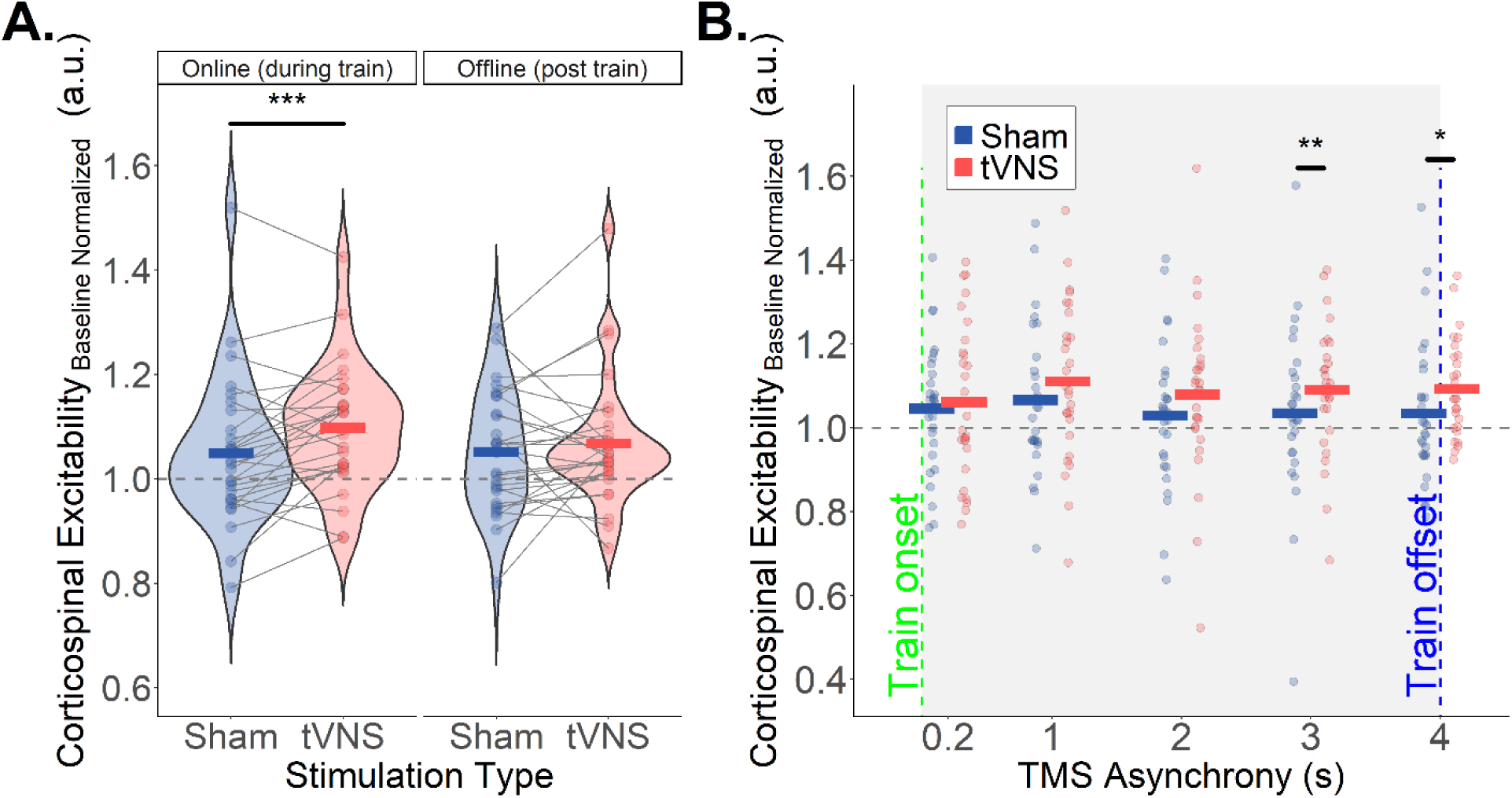
Effect of tVNS on mean corticospinal excitability. **A.** tVNS significantly increased mean corticospinal excitability relative to sham when measured online during stimulation trains, but not offline after stimulation trains. Crossbars represent grand means, individual dots represent participant means, and lines link participants across tVNS and sham conditions. **B.** tVNS-induced increases in corticospinal excitability were primarily driven by measurements taken in the latter half of the stimulation trains (+3 and +4 s post train onset). Crossbars represent grand means, and individual dots represent participant means. \**p*<0.05, \*\**p*<0.01, \*\*\**p*<0.0005.

A follow-up GLMER examining individual timepoints within the train revealed a significant Time × Stimulation Type interaction (β = 0.03, 95% CI [0, 0.03], *p* < 0.05). Excitability was higher in tVNS than sham at +3 s (μ = 0.081 ± 0.03, *p* < 0.01) and +4 s (μ = 0.07 ± 0.03, *p* < 0.05), but not earlier (+0.2, +1, +2 s; all *p* > 0.17; **Figure 2B**). Thus, tVNS online effects were driven by measurements in the latter half of the stimulation train.

### 3.2 tVNS trains drive online increases in pupil size

A cluster-based paired *t*-test revealed greater pupil size during tVNS than sham from 1.1 to 3.2 s after train onset (*p* < 0.05; **Figure 3A**). A GLMER on mean pupil size within this window confirmed a significant Stimulation Type effect (β = 70, 95% CI [35, 104], *p* < 0.001), with higher pupil size during tVNS (μ = 70 ± 17, *p* < 0.0005; **Figure 3B**).

**Figure 3.**
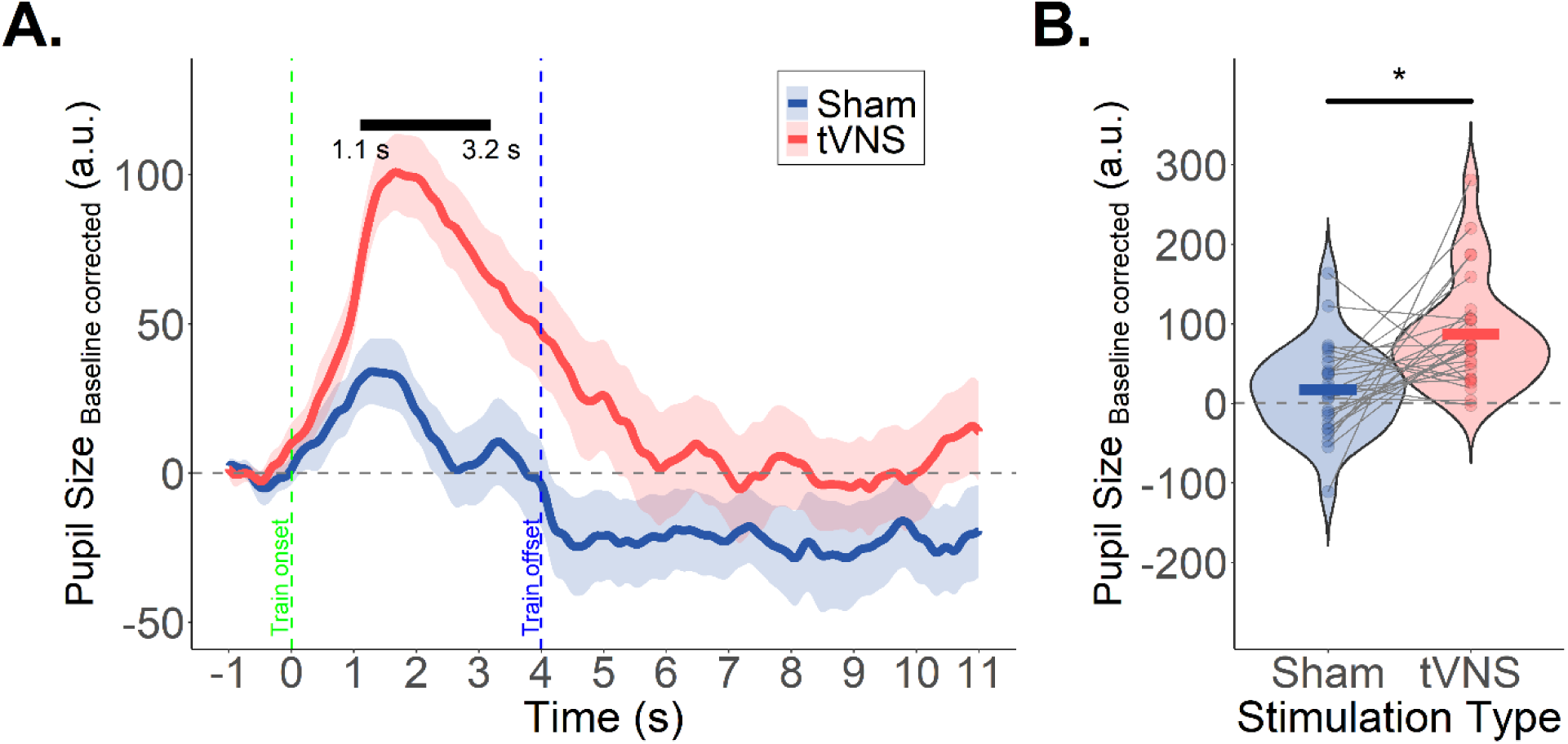
Effect of tVNS on pupil size. **A.** Grand mean of pupil size measurements corrected for pre stimulation train onset pupil size. Shaded regions represent the standard error of the grand mean. tVNS significantly increased pupil size relative to sham from 1.1 seconds post train onset to 3.2 seconds post train onset (black bar). **B.** Mean pupil size was significantly greater under tVNS compared to sham within the significant window outlined in A. Crossbars represent grand means, individual dots represent participant means, and lines link participants across tVNS and sham conditions. \**p*<0.0005.

### 3.3 tVNS-related changes in corticospinal excitability and pupil size are uncorrelated

An LMER showed no significant relationship between tVNS-induced changes in pupil size and corticospinal excitability (β = 30, 95% CI [−190, 260], *p* = 0.79) at any timepoint where MEPs were elicited.

### 3.4 TMS pulses drive increases in pupil size

A cluster-based paired *t*-test showed greater pupil size after TMS than noTMS from 0.47 to 4.3 s post-pulse (*p* < 0.05; **Figure 4A**). A GLMER on mean pupil size within this window confirmed a significant Stimulation Type effect (β = 79, 95% CI [42, 116], *p* < 0.001), with larger pupil size during TMS trials (μ = 79 ± 18, *p* < 0.0005; **Figure 4B**).

**Figure 4.**
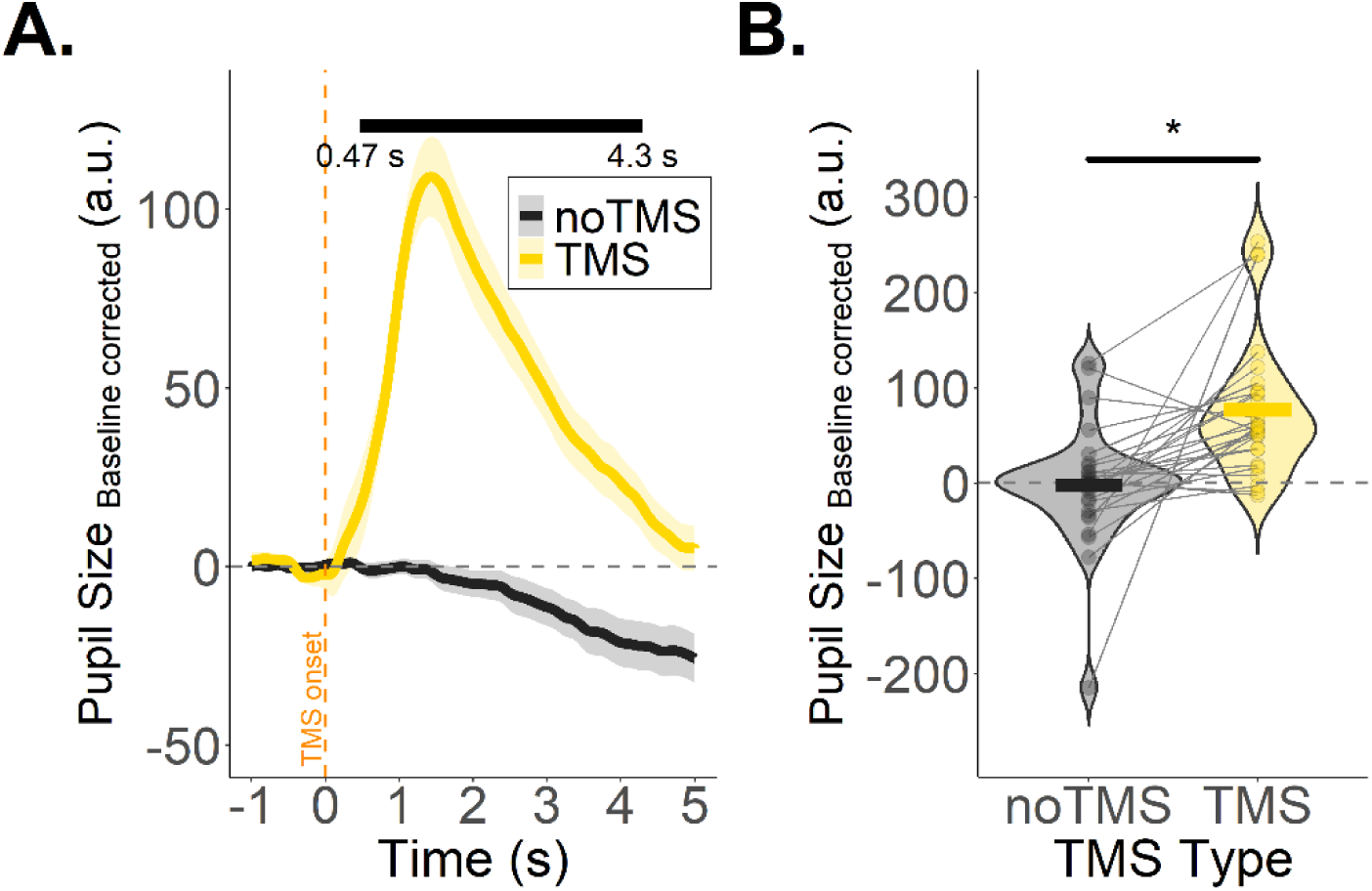
Effect of TMS on pupil size. **A.** Grand mean of pupil size measurements corrected for pre TMS pulse pupil size. Shaded regions represent the standard error of the grand mean. TMS significantly increased pupil size relative to matched trials with no TMS from 0.47 seconds post TMS onset to 4.3 seconds post TMS onset (black bar). **B.** Mean pupil size was significantly greater after TMS compared to matched trials with no TMS within the significant window outlined in A. Crossbars represent grand means, individual dots represent participant means, and lines link participants across tVNS and sham conditions. \**p*<0.0005.

## 4 Discussion

The current study demonstrates that tVNS drives increases in corticospinal excitability. Importantly, a significant effect of tVNS was found only when corticospinal excitability was measured online during tVNS trains and not offline shortly after tVNS train offset (**Figure 2A**). In line with expectations^7–9^, a significant effect of tVNS on pupil size was also found (**Figure 3**). Average changes in corticospinal excitability during tVNS, relative to sham, did not relate to tVNS-induced changes in pupil size. Interestingly, TMS pulses were also found to increase pupil size (**Figure 4**), as has been reported in previous work^63,64^. The degree of increase in pupil size after TMS was similar to that observed during tVNS trains. This suggests that TMS pulses may be capable of inducing NE release, although the precise mechanism of effect likely differs from that seen during tVNS. Taken together, these findings deepen our understanding of LC-corticospinal circuits, and open new avenues for investigation in basic and clinical neuroscience.

### 4.1 tVNS drives increases in corticospinal excitability

Previous investigations into the relationship between tVNS and corticospinal excitability have focused on assessing whether corticospinal excitability is altered by a 30-60 minute bout of continuous tVNS. This work failed to find a consistent effect of tVNS on corticospinal excitability^48–50^. This seems surprising, given that excitation of the LC is associated with the release of the excitatory neurotransmitter NE widely throughout the CNS, including motor areas^13–15^. Based on the finding that the effects of tVNS on pupil size are very fast acting and also decay rapidly after tVNS offset, we reasoned that eliciting MEPs after, rather than during, tVNS may be masking the effect on corticospinal excitability. We addressed this by eliciting MEPs “online” at regularly spaced timepoints during repeated tVNS trains and contrasted their amplitude with those elicited “offline” shortly after the offset of tVNS trains. Using this novel approach, we discovered that tVNS drove up to 8% average increases in corticospinal excitability relative to sham, but only when measured online and not offline (**Figure 2**).

### 4.2 Pupil size is increased by tVNS but changes are uncorrelated with corticospinal excitability

The effect of tVNS on pupil size largely mirrored what has previously been reported^7–9^. Average pupil size was significantly greater under tVNS compared to sham from 1.1 seconds to 3.2 seconds post train onset (**Figure 3**). Average pupil size under tVNS peaked at ∼1.5 seconds post tVNS onset and was more than double that observed under sham within the time window of significant difference. Importantly, sham was matched for perception of tVNS intensity for each participant. Therefore, while sensory inputs are known to induce changes in pupil size^65^ as is evidenced in our data by the small increase in pupil size under sham, the much larger increase under tVNS cannot be driven by sensory input alone and is likely driven by LC activation.

Interestingly, an exploratory analysis aimed at assessing the degree to which changes in pupil size and corticospinal excitability under tVNS were related revealed no significant relationship between the two measures. This is further reflected in the different temporal profiles of the two measures. While pupil size was shown to ramp rapidly towards a peak at around 1.5 seconds post stimulation onset and then gradually decay, corticospinal excitability was shown to build more slowly across time, becoming significantly greater under tVNS compared to sham only at 3- and 4-seconds post stimulation train onset.

When considering the distinct anatomical circuitry involved in modulating both neurophysiological measures, an absence of correlation may not be surprising. The effect of LC activation on pupil size is thought to be driven by concurrent inhibition of the Edinger–Westphal nucleus and excitation of the intermediolateral cell column^66,67^, which synapse directly with the ciliary ganglion and superior cervical sympathetic ganglion respectively. These nuclei in turn directly innervate the iris sphincter and ciliary muscles and exert control over pupil dilation. The precise pathway connecting LC with corticospinal neurons is not known, but it is likely to be far more nebulous than that observed between LC and the iris. The LC does have efferent projections to M1^66,68,69^ which could be the primary pathway for the excitation of corticospinal neurons observed in the current study. However, the LC also has efferent projections to a multitude of other cortical and subcortical regions, including the somatosensory cortices^68,69^, the parietal cortices^70^, the prefrontal cortices^70^, the thalamus^71,72^, the cerebellum^73^, and the spinal cord^74^, which themselves also have known direct and/or indirect connectivity with corticospinal neurons^17,75–82^. Therefore, the changes observed in corticospinal excitability under tVNS are likely driven by the operation of multiple cortical circuits in parallel. While many more studies are needed to pin down the precise anatomical pathway underpinning the increase in corticospinal excitability observed under tVNS, an appreciation of the fundamental difference in the complexity underpinning the MEP effect versus that observed in pupil size may help to explain the lack of correlation between both measures in the current study.

### 4.3 TMS pulses produced similar increases in pupil size as tVNS

While single pulse TMS is primarily used to elicit MEPs to provide a proxy measure for corticospinal excitability, previous studies have also shown that TMS pulses can drive increases in pupil size^63,64^. Therefore, we analyzed the effect of TMS pulses on pupil size in the current study to examine how they compare with changes induced by tVNS. Only TMS pulses that were delivered after the offset of sham stimulation were included to avoid any contaminating effects of tVNS on the pupil signal. Interestingly, TMS pulses were found to increase pupil size to a similar degree as tVNS trains (**Figure 4**). The precise mechanism through which TMS pulses increase pupil size is not well understood. One possibility is that TMS pulses over M1 directly activate the LC, which in turn initiates the pupil response. However, this seems unlikely as although the cortex does have some descending efferent projections to the LC, these are located in the prefrontal^83^ and anterior cingulate cortices^84^, with no evidence for direct projections from the sensorimotor cortices. Another more likely possibility is that the sensorial effects of TMS pulses indirectly increase LC activity via sensory relays. Sensory events are known to drive phasic increases in pupil size^65^, particularly highly salient^85^ and multisensory events^51,86–88^. This is particularly relevant to TMS pulses, which evoke near simultaneous sensory processing of (1) the sensation of the pulse on the scalp, (2) the sound of the pulse, and (3) the sensation of the MEP. In this sense, the increases in pupil size in the wake of TMS pulses could be thought of as more potent version of the small change in pupil size observed after sham stimulation to the earlobes, driven by the additional multisensory features of the stimulus.

### 4.4 Future directions

Results from the current study have broader implications for the use of tVNS in future basic and clinical research. From a practical point of view, they support the use of short train-based approaches where tVNS is applied at specific timepoints within a trial rather than before or constantly throughout a block of trials, as this may reduce the capacity for the brain to compensate for the effects of tVNS^52–54^. They also indicate that 4 seconds is a suitable train length for inducing effects in M1, and that TMS pulses should be timed near the end of the train to best capture the effects of tVNS on corticospinal excitability.

In more conceptual terms, the finding that tVNS drives increases in corticospinal excitability opens the door for use of tVNS within behavioral paradigms where changes in behavioral state are underpinned by modulation of corticospinal excitability. Indeed, while we found relatively modest average effects of tVNS on corticospinal excitability (8% increase), effects may be larger during behavior when the LC is endogenously engaged. For example, corticospinal excitability is reduced during preparation of simple movements^18–28,31–33^, and the application of tVNS during preparation may help delineate the role of the LC in this so-called preparatory suppression^16–18^. Furthermore, the transition from preparation to movement after the presentation of an imperative “Go” stimulus is also associated with a rapid increase in corticospinal excitability in the selected effector^24,25,30^ and a concurrent rapid decrease in non-selected effectors^29^, the onset of which may represent an underlying shift in the neural dynamics of M1 between preparatory and execution related states^30^. The addition of tVNS in paradigms focused on this effect may help clarify the precise role of the LC in this transition.

A particularly interesting overarching question for future studies combining tVNS, TMS, and behavior will be whether the effects of tVNS are always excitatory or if they depend on behavioral state. Within the domain of preparatory suppression, for example, we might expect the application of tVNS to instead *strengthen* suppressive effects, since greater suppression is associated with faster responses^22,32^ and therefore greater task success, and the LC is broadly thought to intervene to maximize task success within reward rich environments^89,90^. Even if NE (the output of the LC) is an excitatory transmitter, it could achieve net suppression of corticospinal neurons through excitation of local inhibitory interneurons.

Our results also have implications for clinical research. tVNS has been touted as treatment for motor system diseases including stroke^91^, Parkinson’s disease^92^ and multiple sclerosis^93^, since it is believed to have the potential to broadly excite the brain including the corticospinal tract, which is often damaged or dysfunctional in these populations^41–46^. Many existing studies have been interested in the potential “priming” effects of tVNS, with the belief that a bout of tVNS would place the brain in an excited and plastic state which could be taken advantage of to maximize neural recovery in a following behavioural therapy session. This is reflected in existing literature examining the effects of tVNS on corticospinal excitability, where measures of corticospinal excitability were taken before and after a 30-60 minute bout of continuous tVNS^48–51^. Our finding that corticospinal excitability was only found to be reliably affected by tVNS “online” while stimulation is actively applied speak against the notion of tVNS priming. Instead, they support an approach in which tVNS is dynamically applied concurrently with behavior, possibly in short bursts at salient timepoints within a given trial. Indeed, by using such an approach previous investigations in our lab have been able to identify significant effects of tVNS on decision making accuracy^7,55^. Furthermore, this approach has already been demonstrated to have superior efficacy in clinical studies assessing the effects of invasive VNS in chronic stroke populations^94^. Therefore, assessing the efficacy of a paired tVNS/behaviour approach in clinical populations and identifying the optimal timing, duration, intensity and dose of such tVNS bursts represent important questions for future work.

### 4.5 Conclusions

The current study demonstrates that the increases in pupil size at rest observed during tVNS are also observed in corticospinal excitability. The effect of tVNS on corticospinal excitability only emerges during active stimulation. The temporal profile of change in corticospinal excitability differs from that observed for pupil size, with the greatest increases observed in the late rather than early period of stimulation. This is likely driven by differences in the anatomical circuits underpinning each effect, and future studies leveraging other TMS techniques such as dual coil TMS, directional TMS, and paired pulse TMS along with functional neuroimaging will help to better define the precise pathway of the effect of tVNS on corticospinal excitability. Evidence that corticospinal excitability is modulated by tVNS opens new avenues for applying the method to further understanding of LC-corticospinal circuits during various behavioral paradigms, including motor preparation and decision making. In terms of future clinical applications, given that our results indicate that the effects of tVNS on cortical activity are strongest during active stimulation, they support the approach of pairing tVNS with behavioural interventions to maximize clinical outcomes.

## CRediT authorship contribution statement

**Ronan Denyer:** Conceptualization, Methodology, Software, Validation, Formal analysis, Investigation, Data curation, Writing – original draft, Writing – review & editing, Visualization, Project administration, Funding acquisition. **Su Shiyong:** Methodology, Software, Formal analysis, Writing - Review & Editing. **Mantosh Patnaik:** Investigation, Writing - Review & Editing **Julie Duque:** Conceptualization, Methodology, Validation, Resources, Writing – review & editing, Project administration, Funding acquisition.

## Acknowledgement

This work was supported by a grant awarded to RD by the Belgian National Funds for Scientific Research (FRS-FNRS:FC 59664), by a Coordinated Research Project (ARC) grant awarded to JD by UCLouvain (Coaction), and by research credits awarded to JD by FNRS (PDR UrgeToAct: T007023F and CDR-AROUSAT: J007424F).

## Data Availability Statement

The data that supports the findings of this study are currently available on request, and will be placed in an open repository upon publication of the manuscript by a peer-reviewed journal.

